# Non-linear frequency-dependence of neurovascular coupling in the cerebellar cortex implies vasodilation-vasoconstriction competition

**DOI:** 10.1101/2021.07.30.454430

**Authors:** Giuseppe Gagliano, Anita Monteverdi, Stefano Casali, Umberto Laforenza, Claudia A.M. Gandini Wheeler-Kingshott, Egidio D’Angelo, Lisa Mapelli

## Abstract

Neurovascular coupling (NVC) is the process associating local cerebral blood flow (CBF) to neuronal activity (NA). Although NVC provides the basis for the blood-oxygen-level-dependent (BOLD) effect used in functional MRI (fMRI), the relationship between NVC and NA is still unclear. Since recent studies reported cerebellar non-linearities in BOLD signals during motor tasks execution, we investigated the NVC/NA relationship using a range of input frequencies in acute mouse cerebellar slices of vermis and hemisphere. The capillary diameter increased in response to mossy fiber activation in the 6-300Hz range, with a marked inflection around 50Hz (vermis) and 100Hz (hemisphere). The corresponding NA was recorded using high-density multi-electrode arrays and correlated to capillary dynamics through a computational model dissecting the main components of granular layer activity. Here, NVC is known to involve a balance between the NMDAR-NO pathway driving vasodilation and the mGluRs-20HETE pathway driving vasoconstriction. Simulations showed that the NMDAR-mediated component of NA was sufficient to explain the time-course of the capillary dilation but not its non-linear frequency-dependence, suggesting that the mGluRs-20HETE pathway plays a role at intermediate frequencies. These parallel control pathways imply a *vasodilation-vasoconstriction competition hypothesis* that could adapt local hemodynamics at the microscale bearing implications for fMRI signals interpretation.

## Introduction

The neurovascular coupling (NVC) comprises mechanisms that link neuronal activity (NA) and vascular motility, adjusting the cerebral blood flow (CBF) to the energy demand of neural tissue. In the last couple of decades, increasing attention has been devoted to addressing NVC mechanisms, both in physiological and pathological conditions. The *metabolic hypothesis*, with the rapid glucose and oxygen consumption by active neural cells driving blood vessel responses, has long been considered the basis for the interpretation of blood oxygen level-dependent (BOLD) signals recorded using functional magnetic resonance imaging (fMRI). However, the cellular and molecular mechanisms involved have not yet been clarified ^1-4^, and the validity of this hypothesis was undermined by several observations where CBF was not directly linked to changes in oxygen or glucose concentration ^5-7^. Further studies supported the *neurogenic hypothesis*, in which CBF changes are determined by direct signaling through molecules released by neurons or glial cells ^8-10^. Interestingly, the role of astrocytes in determining the vascular tone has been emphasized in the *set-point hypothesis*, balancing dilation and constriction to maximize vessel responses to NA ^11^. Additional mechanisms as intrinsic vascular properties are involved as well ^12^ and might contribute to region-specific differences ^13^. Finally, differences in vasodilation and vasoconstriction agents have been described throughout the brain ^14^. It is much likely that all these microscale mechanisms intervene to some extent in determining the CBF changes following neuronal activation, probably with different relevance depending on the region involved ^7^, complicating the interpretation of macroscale phenomena like the BOLD signal ^3, 15^.

In the cerebellum, NVC shows particular properties. The densely packed granular layer is the main determinant of changes in cerebellar energy consumption and the granule cells, which are NOergic and release nitric oxide (NO) in both the granular and molecular layers ^16, 17^, are likely to be the primary controllers of NVC ^18-20^. Vasodilation in the granular layer is entirely mediated by neuronal N-methyl-D-aspartate receptor (NMDAR)-dependent production of NO, while the main vasoconstriction pathway is the metabotropic glutamate receptors (mGluRs)-dependent production of 20-hydroxyeicosatetraenoic acid (20-HETE), presumably released by astrocytes ^21^ and blocked by NO itself ^8^. The integration of vasodilation and vasoconstriction signals is likely to occur in pericytes, in agreement with their role in controlling microvessel caliber and initiating the BOLD signal ^22, 23^. A possible NVC region specificity has been recently suggested by the evidence of different BOLD responses in the cerebellar cortex during the execution an intensity-graded grip force motor task in humans, with the most substantial effects in vermis lobule V and hemisphere lobule VI ^24, 25^, but it is unclear whether these differences simply reflect different cortical inputs or local NVC factors. Moreover, reports of non-linear BOLD response alterations due to pathology such as multiple sclerosis make understanding NVC factors very important for explaining the mechanisms of disease ^26, 27^.

Several studies reported a direct correlation between NA and blood perfusion, where NA is measured as the summed local field potentials (∑LFP) evoked during electrical or sensory stimulation ^28-36^. Generally, the stimulus-evoked changes in CBF (or related parameters) linearly correlate with NA at increasing activation rates. Non-linearities have been reported mainly as a *ceiling-effect* due to a saturation of CBF increase at certain frequencies or stimulus amplitudes ^28-30, 37, 38^, usually with increasing perfusion at increasing frequencies. Interestingly, few cases have been reported of perfusion decreasing at higher frequencies, well explained by the corresponding decrease of ∑LFP ^33, 36^. In summary, the studies so far reported a linear correlation of blood perfusion with NA, with some non-linearity mainly due to structural limitations of vessel motility.

Here, we investigated NVC in the granular layer of acute cerebellar slices by electrically activating mossy fibers at 6 to 300 Hz and measuring capillary diameter changes and NA responses in the vermis lobule V and hemisphere lobule VI. NVC increased with frequency but showed a marked inflexion at intermediate frequencies that was not attributable to saturation of the effect. An advanced realistic computational model of the granular layer was used to dissect the NA features best correlating with vessel responses, confirming the key role of NMDAR in determining the time-course of vessel dilation. Unexpectedly, NA-related parameters (including ∑LFP) were not sufficient to explain the specific frequency-dependence profile of cerebellar NVC, suggesting that additional mechanisms might be involved. The hypothesis of a NA-driven frequency-dependent competition between vasodilation and vasoconstriction pathways is therefore proposed as hypothesis and thoroughly discussed.

## Methods

Animal maintenance and experimental procedures were performed according to the international guidelines of the European Union Directive 2010/63/EU on the ethical use of animals and were approved by the local ethical committee of the University of Pavia (Italy) and by the Italian Ministry of Health (authorization n. 645/2017-PR). According to this authorization, the sample size was estimated *a priori* using G*Power software (Wilcoxon-Mann-Whitney test, effect size 1.5, significance level 0.05) yielding an actual Power of 0.82. Reporting complies with the ARRIVE guidelines (Animal Research: Reporting in Vivo Experiments).

### Preparation of acute cerebellar slices

Acute cerebellar slices were obtained from 17-23 days old C57BL/6 mice of both sexes, as reported previously ^21, 39-41^. Briefly, mice were anesthetized with halothane (Aldrich, Milwaukee, WI) and killed by decapitation. Acute parasagittal slices of 220μm thickness were cut from the cerebellar vermis and hemisphere with a vibroslicer (VT1200S, Leica Microsystems). During the cutting procedure, slices were maintained in cold Krebs solution. Then, slices were recovered for at least 1h in the same solution before being incubated for 1h with 75nM U46619 (Abcam), a thromboxane agonist. This procedure is commonly used to restore the original vascular tone in acute slices ^21, 42-44^. For recording, slices were transferred to a 2-ml recording chamber mounted on the stage of an upright microscope (Slicescope, Scientifica Ltd, UK) or to the recording chamber of the HD-MEA (Arena biochip, Biocam X, 3Brain AG). In both cases, Krebs solution was perfused (2 ml/min) and maintained at 37°C with a Peltier feedback device (TC-324B, Warner Instruments, Hamden, CT) for the entire duration of the experiments. Krebs solution for slice cutting and recovery contained (in mM): 120 NaCl, 2 KCl, 1.2 MgSO_4_, 26 NaHCO_3_, 1.2 KH_2_PO_4_, 2 CaCl_2_, and 11 glucose, and was equilibrated with 95% O_2_-5% CO_2_ (pH 7.4). Krebs solution perfused during recordings was added of 75nM U46619. All drugs were obtained from Sigma Aldrich, unless otherwise specified.

### Immunofluorescence staining

Pericytes and capillaries were stained in the granular layer of both cerebellar vermis and hemisphere slices as previously described ^21, 44^, focusing respectively on lobule V and lobule VI. First, slices (220 μm-thick) were fixed with 4% paraformaldehyde in PBS for 25 min in a Petri dish. Secondly, slices were washed three times before being permeabilized and blocked with 5% Triton X-100, 10% BSA added to PBS, overnight at 4°C. Thirdly, samples were incubated for 24 h at 25°C on a rotary shaker with rabbit anti-NG2 chondroitin sulfate proteoglycan (Millipore, cat no. AB5320, 1:200 dilution) and FITC-isolectin B4 (Sigma-Aldrich, cat. no. L2895; 1:200 dilution of a stock solution of 2 mg/ml FITC-IB4 in PBS) primary antibodies to stain respectively pericytes and blood vessels. Before the incubation with secondary antibodies, slices were washed three times (each of 15 min) with PBS. Fourthly, samples were incubated for 4 h at 25°C with fluorescent secondary antibody Rhodamine Red-X-AffiniPure Goat Anti-Rabbit IgG (Jackson ImmunoResearch Inc. cat. no.111-295-045; 1:500 dilution). Finally, slices were washed three times with PBS and mounted on miscroscope slides, ProLong® Gold antifade reagent with DAPI (Molecular Probes) and coverslips affixed. Fluorescence of cerebellar samples was observed with a TCS SP5 II LEICA confocal microscopy system (Leica Microsystems, Italy) furnished with a LEICA DM IRBE inverted microscope. The confocal system exploited a 20X, 40X, and 63X objectives to acquire images. All acquisition files were visualized by LAS AF Lite software (Leica Application Suite Advanced Fluorescence Lite version 2.6.0) installed on a desktop PC. Negative controls were carried out in parallel by treating slices with non-immune serum during the incubation procedures.

### Time-lapse acquisition and analysis of capillary diameter changes

Granular layer vessels were identified using bright-field microscopy in parasagittal slices of cerebellar vermis and hemisphere. Vermis lobule V and hemisphere lobule VI were first identified using a 4X objective (XL Fluor 4X/340, N.A.: 0.28, Olympus, Japan). Secondly, the granular layer was inspected using a 60X objective (LumPlanFl 60X/0.90 W, Olympus, Japan) which allowed to locate capillaries and pericytes. Only vessels with an inner diameter <10μm and surrounded by at least 1 pericyte (arrowheads in **Fig.1B**) were considered. Only one capillary per slice was used for experiments. In *ex-vivo* conditions, capillaries lose intraluminal flow and pressure due to the mechanical stress of the slicing procedure. Slices were incubated with 75nM U46619, a thromboxane agonist that re-establish the vascular diameter mimicking physiological conditions, prior to the experiment, as previously described ^21^. Once the focus of the objective was adjusted on the capillary walls, the pericyte soma could no longer be clearly visible. With their long projections, pericytes regulate the caliber of distant portions of the vessels ^23^. Therefore, pericytes membrane adjacent vessel walls may appear blurry when the focal plane was chosen.

Mossy fibers were stimulated with 15V stimuli at 6Hz, 20Hz, 50Hz, 100Hz, and 300Hz for 35s, using a bipolar tungsten electrode (Warner Instruments, UK) in cerebellar lobule V or VI in vermis and hemisphere slices, respectively. The experiments were performed in Krebs solution perfused (2 ml/min) using a peristaltic pump (ISMATEC) and maintained at 37°C with a Peltier feedback temperature controller (TC-324B; Warner Instrument Corporation). In all cases, mossy fibers stimulation induced a significant vasodilation, as showed in **Fig.1B**. Capillary responses in the granular layer were detected about 200μm distant from the stimulating electrode (average distance in vermis slices: 131.93±5.60μm; n=50 and hemisphere slices: 135.45±5.28μm; n=49; vermis vs hemisphere p=0.648). The time-lapse bright-field images of capillaries caliber changes were obtained using a CCD camera (DMK41BU, Imaging Source, Germany), controlled by the IC-capture 2.1 software (Imaging Source, Germany) to acquire 1 image every second (before, during and after mossy fibers stimulation) with 5ms exposure time. Image sequences were analyzed offline using the ImageJ software, as previously described ^21^. The portion of the vessel showing the maximal effect was considered for the analysis. Data were compared using statistical paired and unpaired Student’s *t* test and with ANOVA test. First, we applied the normality test to check the normal distribution of data. Then, the parametric one-way ANOVA and finally the Fisher *post-hoc* tests were used to validate the statistical significance. Data were considered statistically significant with p<0.05 and are reported as mean ± SEM (standard error of the mean).

### Electrophysiological recordings of neuronal activity in cerebellar slices

Neuronal activity was recorded as local field potential (LFP) in the granular layer of vermis lobule V and hemisphere lobule VI during mossy fibers stimulation at 37° C. Slices were treated as for the time-lapse imaging and placed on the glass recording chamber of a high-density multi-electrode array (HD-MEA) system (BioCAM X, 3Brain AG, Switzerland). The recording probe was a CMOS biochip with 4096 recording microelectrodes arranged in a 64 x 64 matrix, covering an area of 2.67mm x 2.67mm (Arena biochip, 3Brain AG). The electrodes size was 21μm x 21μm with a pitch of 42μm. Neuronal activity was sampled at 17840.7 Hz/electrode and acquired with BrainWave X software (3Brain AG). Krebs solution was continuously perfused during the experiment using a peristaltic pump (ISMATEC) at the rate of 2ml/min and maintained at 37°C with a Peltier feedback temperature controller (TC-324B; Warner Instrument Corporation). Stimulation was provided by a tungsten bipolar electrode (Warner Instruments) placed over the mossy fibers using a micromanipulator (Patch-Star, Scientifica Ltd). The stimulator unit was embedded in the Biocam X hardware and set to deliver current pulses of 50µA for 200µs duration, matching the same stimulation conditions used in time-lapse imaging recordings of capillary responses. Due to technical limitations, the 300Hz stimulation frequency was not included in the analysis. In the granular layer, neuronal response to mossy fiber stimulation originated LFPs. In both vermis and hemisphere, the LFP shows a typical N_1_–N_2a_-N_2b_–P_2_ complex: N_1_ corresponds to presynaptic volley activation, N_2a_-N_2b_ are informative of granular cells synaptic activation and P_2_ is likely to represent currents returning from the molecular layer ^45^. In order to characterize granule cells responses to stimulation, the analysis was focused on N_2a_ and N_2b_ peaks amplitude and time to peak. Recordings were exported and analyzed with Matlab (Mathworks). Given the biochip electrodes impedance of 1-5MΩ, the electrode surface of 20µm, and the densely packed granule cells around, a good estimate of the number of granule cells contributing to the LFP in each channel is (at least) 10-15 cells. To match the distances used for capillary responses, the signals recorded in channels located beyond 200µm from the stimulation electrode were not included in the analysis. Given the high spatial resolution of the HD-MEA used, the granular layer responses in a 200µm range were detected by nine channels per slice. The average distance from the stimulating electrode of the electrophysiological responses was 133 ± 3.32 μm (n=20 slices; n=159 electrodes), not statistically different from the capillary average distances from the stimulating electrode in time-lapse imaging (133±3.83 μm, n=99 vessels; unpaired Student’s t test p=0.99). The cumulative LFP during the stimulation was calculated to compare the NA trend to the capillary responses in the same conditions.

### Computational model of the granular layer

Simulations have been performed using a detailed computational model of the granular layer (GL) network ^46^ endowed with biophysically realistic single cell models of granule cells (GrCs) and Golgi cells (GoCs), as well as a simplified reconstruction of mossy fibers terminals, the glomeruli (gloms). Detailed reconstruction of synaptic dynamics and receptors was also included, allowing to reproduce the effect of α-amino-3-hydroxy-5-methyl-4-isoxazolepropionic acid (AMPA), NMDA, and gamma-aminobutyric acid (GABA) receptors mediated currents. The full GL network had a volume of 800 x 800 x 150 µm^3^; circuit organization and connectivity has been reproduced according to specific connectivity rules, accounting for both geometric and statistical data (see also ^47^ for a similar approach). The original version of the granular layer network faithfully reproduced the cell-density distributions experimentally observed in the cerebellar vermis: the full network included 384.000 GrCs, 914 GoCs and 29.415 gloms. In order to reconstruct a network of the cerebellar hemisphere, different density distributions had been considered: cell data coming from the Allen Brain Atlas (mouse data, https://mouse.brain-map.org/) for the Crus (i.e., the hemisphere) have been collected, analyzed, and compared to those coming from the Lingula (i.e., the vermis). A comparative analysis showed that GrCs and GoCs densities in the hemisphere were respectively 64% and 89% higher than in the vermis. Finally, gloms density has been estimated assuming that the GrCs / gloms ratio should be kept constant. This led to a network reconstruction of the hemisphere-GL with 630.419 GrCs, 1.733 GoCs and 48.291 gloms. It should also be noted that the GrCs / GoCs ratio changed significantly from the vermis (∼ 420:1) to the hemisphere (∼ 364:1).

The experimental protocol has been reproduced with the network model as follows: a subset of gloms was stimulated with continuous trains at five different frequencies, 6, 20, 50, 100, and 300 Hz, 35 seconds per simulation. Gloms were selected according to a simple geometrical rule (see also ^46^): all the gloms included within a sphere centered in the middle of the GL network and with radius equal to 27.7µm were stimulated with the same train. Given the different cell-densities, the total amount of excited GrCs was 1290 and 1996 for vermis and hemisphere networks, respectively. Notice that the number of GrCs contributing to the simulated response is similar to the estimated total number of GrCs contributing to the LFP signal (see above: 10-15 neurons per electrode, 9 electrodes per slice, 10 slices per condition, resulting in a number of neurons contributing to the total average LFP above 900-1350 units).

NMDAR-mediated currents were recorded for each stimulated GrC. In order to compare NO production to NMDA currents over time, the cumulative sum of NMDA-current, averaged over each cell, was calculated at different time points: 2, 20, and 35 seconds of the stimulation period. The computational model of GL network has been developed in the NEURON-Python simulation environment; simulations were performed on a 60 cores local cluster (five blades with two Intel Xeon X560 and 24 Gigabyte of DDR3 ram per blade, twelve cores per blade).

### Model validation

The model was validated against the *N*_*2a*_ component of the LFP using the following procedure. The average membrane depolarization of all the active GrCs was computed in the 1ms time-window following the stimuli. This corresponded to the N_2a_ peak observed in the experiments. The procedure was repeated for all stimulation frequency used in the experiments. The resulting set of measurements was normalized and low-pass filtered (Butterworth, 0.2 Hz, 2nd order). These simulated data were compared to experimental data obtained measuring the N_2a_ peaks.

## Results

In the cerebellum, CBF changes mostly rely on capillary diameter changes in the granular layer following change in NA ^21, 22^. Synaptic activation determines the release of vasoactive molecules acting on pericytes, contractile cells that enwrap the capillary wall ^48, 49^. Here, we have analyzed the effect of different input frequencies on NVC in the granular layer of vermis lobule V and hemisphere lobule VI of the cerebellar cortex.

### Anatomical organization of neurovascular components in the granular layer of vermis and hemisphere

In the granular layer of both vermis and hemisphere, the microvessels wall consisted of tightly connected endothelial cells surrounded by granule cells (asterisks in Fig.1) but not by smooth muscle cells, in agreement with previous reports in rats ^21^. Notably, granule cells are in close proximity with pericytes, making the latter very suitable to receive chemical transmitters from activated neurons ^50^. Both in vermis and hemisphere, cerebellar slices showed granular layer capillaries labeled by anti-isolectin B4 and anti-proteoglycan NG2 primary antibodies, which stained vessel and pericyte membranes, respectively ^21, 44^ (**Fig.1A**). The pericytes appeared to enwrap capillaries (arrowheads in **Fig.1A and 1B**), whose pre-contracted average internal diameter was not statistically different (unpaired Student’s *t* test, p=0.13) in the vermis (2.75±0.15μm, n=50) and hemisphere (2.47±0.11μm, n=49). Thus, cerebellar vermis and hemisphere showed similar granular layer organization of neurovascular components, comprised of capillaries, pericytes and granule cells.

### Non-linear frequency-dependent dilation of granular layer capillaries (NVC)

NVC was assessed by measuring capillary diameter changes following mossy fibers stimulation. Mossy fibers stimulation determined a significant vasodilation in granular layer capillaries, both in the vermis and hemisphere, at all the frequency tested (6, 20, 50, 100, 300 Hz; **Supplementary Table 1**) but the amount of dilation varied (**Figs 1C and 2**). In the vermis (**Figs 1C and 2A**), significant differences were evident at 2s (20 Hz vs. 50 Hz), 20s (20 Hz vs. 50 Hz) and 35s (50 Hz vs. 100Hz; 6 Hz vs. 20 Hz, 100 Hz, 300 Hz). In the hemisphere (**Figs 1C and 2A**), significant differences were evident at 2s (50 Hz vs. 300Hz) and 35s (100 Hz vs. 20 Hz, 300 Hz). Consequently, the maximum dilation did not increase linearly with frequency, as illustrated for the points taken at 35s **(Fig.2B)** (the subsequent analysis will focus on this time-point).

**Fig. 1.**
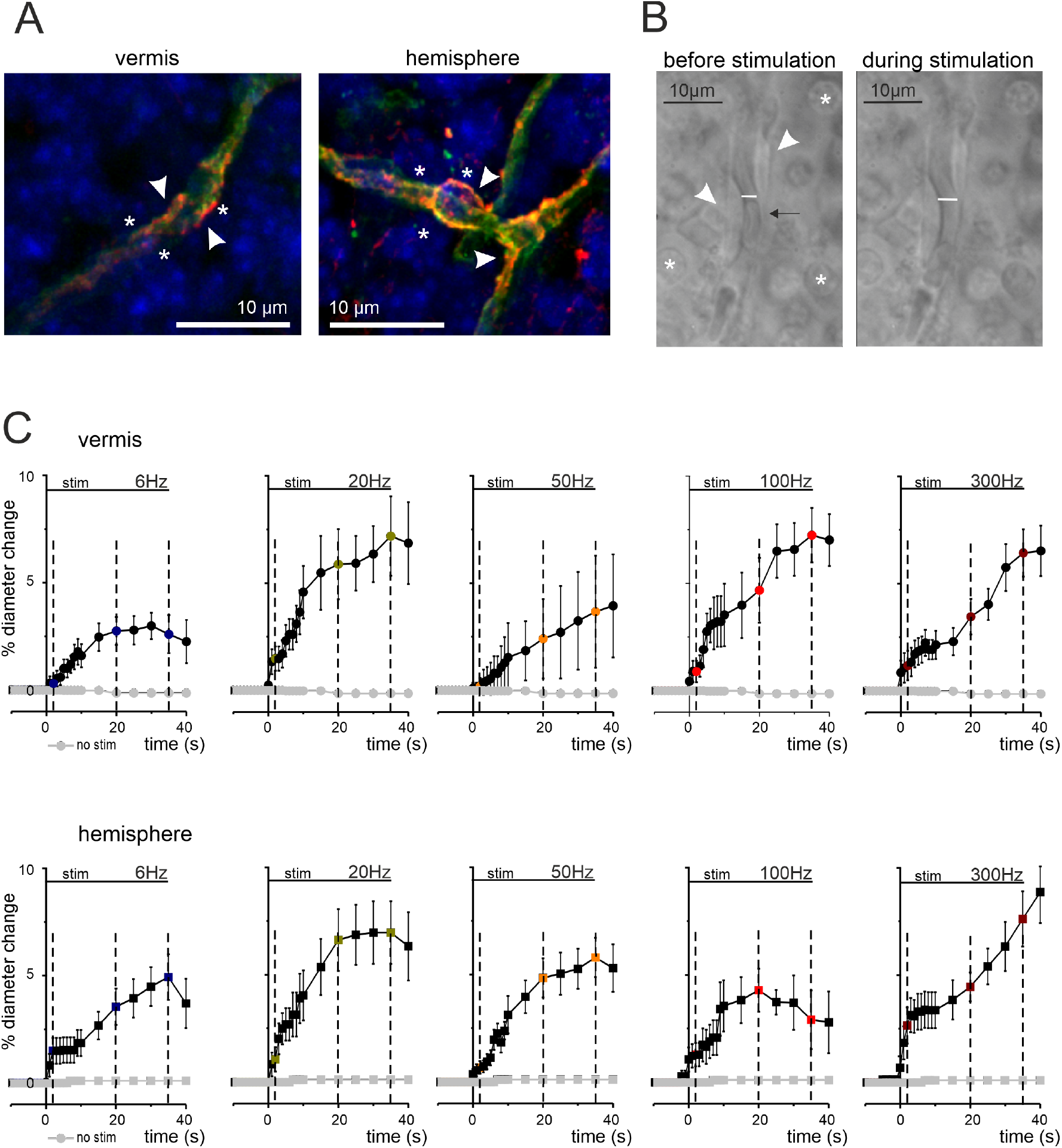
Anatomo-physiological correlates of NVC in the cerebellar granular layer. **A)** Confocal fluorescent images of cerebellar slices stained for IB4 (green), NG2 (red), and DAPI (blue) to identify capillaries, pericytes, and cell nuclei, respectively. In both vermis and hemisphere, granule cells (asterisks) are in proximity of pericytes (arrowheads) located on capillary endothelium (green). **B)** Bright-field images taken before and during mossy fiber stimulations at 50 Hz (vermis). The pericytes (arrowheads) are visible near granule cells (asterisks) and the capillary walls can be easily identified. The internal diameter changes in response to stimulation (white bar). The arrow indicates a red blood cell that moves inside the capillary as a consequence of blood vessel motility. **C)** Average time courses of capillaries dilation during mossy fibers activation at different frequencies in vermis and hemisphere. The figure shows the average percent change in capillaries diameter size in the different conditions tested for the vermis lobule V and hemisphere lobule VI. Mossy fibers were stimulated for 35s at 6, 20, 50, 100, and 300 Hz (stim bar). The dashed lines indicate the time points at 2, 20, and 35s, used to analyze dilation in the subsequent analysis. In each panel, the dilation at the end of stimulus was significantly different compared to pre-stimulus baseline (see Table 1in Supplemental Material). In both panel sets, grey dots represent stability recordings in which stimulation was not delivered (n=8).

**Fig. 2.**
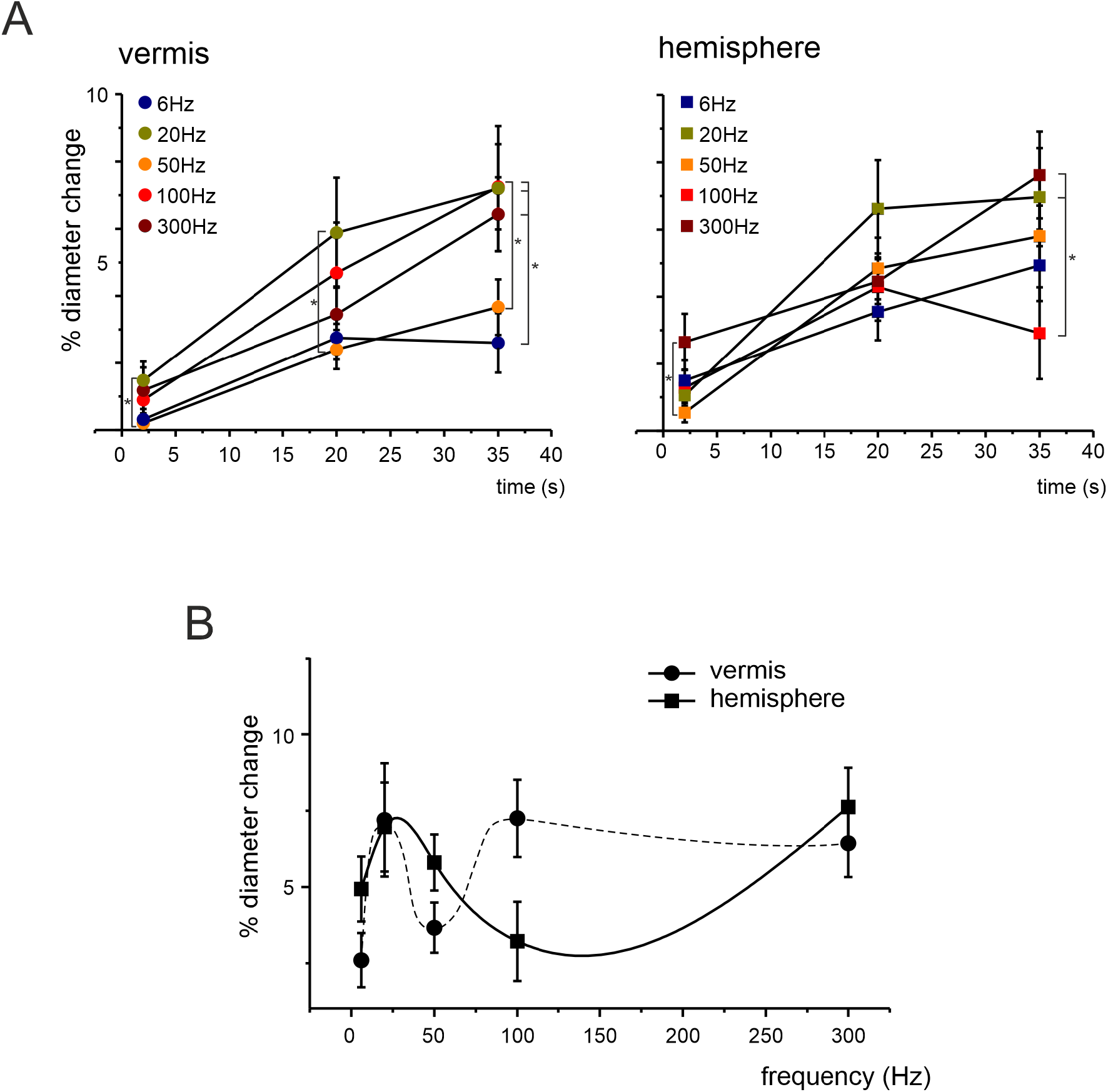
NVC frequency-dependence in the cerebellar granular layer. **A)** Average time course of vasodilation during mossy fibers stimulation at 2, 20, and 35s in capillaries of the vermis and hemisphere. Asterisks indicate pairs of points that were statistically different (* p<0.05; ** p<0.01). Vermis: 2s, 20Hz vs. 50Hz (p=0.039); 20s, 20Hz vs. 50Hz (p=0.0358); 35s, 6Hz vs. 100Hz (p=0.014), 50Hz vs. 100Hz (p=0.049), 6Hz vs. 20Hz (p=0.015), 6Hz vs. 300Hz (p=0.036) (one-way ANOVA; for increasing frequencies n = 9, 10, 10, 10, 11, respectively). Hemisphere: 2s, 50Hz vs. 300Hz (p=0.029); 35s, 100Hz vs. 20Hz (p=0.024), 100Hz vs. 300Hz (p=0.009) (one-way ANOVA; for increasing frequencies n= 9, 10, 10, 10, 10, respectively). **B)** Average dilations at 35s at each frequency tested in the vermis and hemisphere.

The dilation at different stimulation frequencies reached its maximum around 20Hz but showed a decrease at 50-100Hz before attaining maximum dilation at higher frequencies. The maximum dilation in vermis and hemisphere capillaries did not show the same frequency-dependent trend. E.g., at 100Hz, dilation in the vermis was at its maximum (7.25±1.27%, n=10), while in the hemisphere it was around minimum values (3.21±1.29%, n=10, p=0.032). It should be noted that the maximum dilation was similar in the vermis and hemisphere, despite differences in the frequency-dependence profile.

### Granular layer responses to mossy fibers stimulation (NA)

In order to correlate NA with blood vessels dilation, several studies in different brain areas have used local field potential (LFP) cumulative amplitude ^28-36^. Here, the LFPs elicited in the granular layer by mossy fibers stimulation were measured using a high-density multielectrode array (HD-MEA). This technique allowed us to assess the spatial distribution of NA and to record the LFPs at a distance from the stimulating electrode comparable to that of capillaries. In this way, NA was recorded by the same tissue volume that was likely to generate the bulk of the signals that correlate NA to NVC (see Methods; **Fig.3A**).

**Fig. 3.**
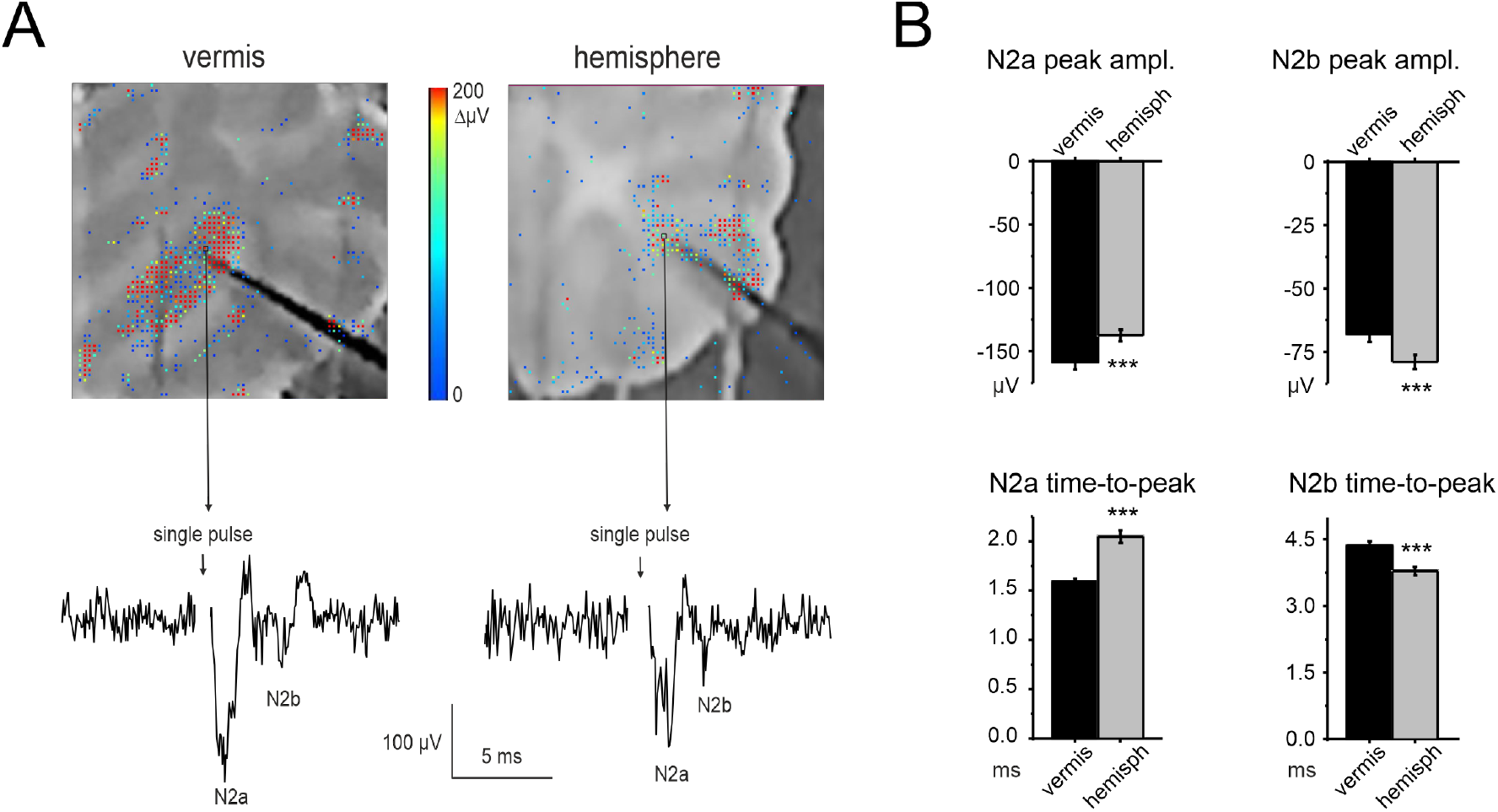
NA recordings in in the cerebellar granular layer. **A)** Example of a vermis and hemisphere cerebellar slices placed on the HD-MEA biochip. The activity following single pulse stimulation of mossy fibers is superimposed in a color-scale (2ms bin). A representative trace is shown from the HD-MEA biochip channel indicated by the arrow. The stimulus intensity and stimulating electrode is the same as used in Fig. 1B. **B)** The histograms show the average peak amplitude and time-to-peak of N_2a_ and N_2b_, both for the vermis and hemisphere (n=10 slices for both). Statistical significance is indicated by asterisks (***p<0.001).

According to previous characterizations of granular layer LFPs ^45, 51^, N_2a_ and N_2b_ peaks derive from synaptic activation of multiple granule cells clustered around the electrode. N_2a_ mainly depends on the AMPAR-mediated component of granule cells responses and on spike synchronicity, while N2b is informative on the NMDAR-mediated component of granule cells responses and its inhibitory control ^45, 52^.

The response to single-pulse mossy fibers stimulation was first characterized in granular layers of the vermis and hemisphere (**Fig.3A**). In the vermis, N_2a_ peaked at 1.59±0.03ms with an amplitude of −158.74±5.53 µV and N_2b_ peaked at 4.36±0.1ms with an amplitude of −67.93±3.04 µV (n=10 slices, 81 electrodes for all measures). In the hemisphere, N_2a_ peaked at 2.05±0.06 ms with an amplitude of −137.39±4.66 µV and N_2b_ peaked at 3.79±0.09 ms with an amplitude of −78.88±2.90 µV (n=10 slices, 76 electrodes for all measures). Thus, N_2a_ had higher peak and shorter time-to-peak in the vermis compared to hemisphere (unpaired Student’s *t* tests, p=0.003 and p=6.85*10^−10^, respectively); conversely, N_2b_ had higher peak and shorter time-to-peak in the hemisphere compared to vermis (unpaired Student’s *t* tests, p=0.01 and p=0.000123, respectively) (**Fig.3B**).

In the same slices, granular layer responses were characterized using the same stimulation patterns previously used to assess vasodilation (except for 300Hz, see Methods). The time-course of N2a peak amplitude at different frequencies for vermis and hemisphere is reported in **Fig.4A**. In all cases, during the stimulation, N_2a_ peak amplitudes decreased and attained a plateau. The decrease was more evident at higher frequencies (see **Supplementary Table 2** for the percent change at the end of the stimulation). N_2b_ peak amplitudes showed a trend similar to N_2a_ (**Fig.4A**). It should be noted that, despite identical stimulation intensity, N_2a_ was usually smaller in hemisphere than vermis (e.g., at 2s, 20s, 35s; n=10 for both, unpaired Student’s t test p<0.05).

**Fig. 4.**
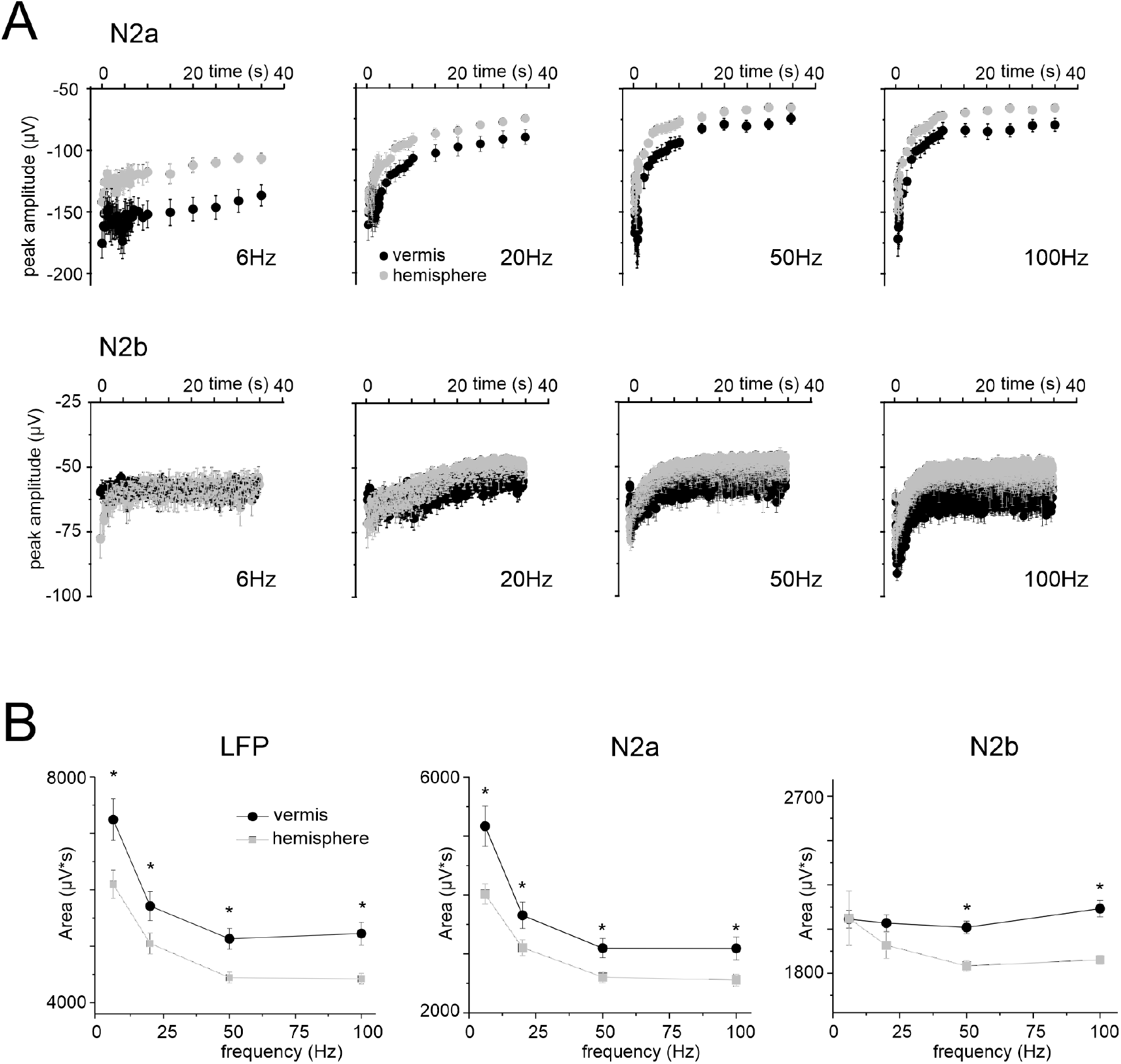
Time- and frequency-dependence of LFPs during stimulation. **A)** The plots show the time course of N_2a_ and N_2b_ peak amplitude in response to stimulation at different frequencies, recorded in 10 vermis and 10 hemisphere slices (the responses in the right part of the N_2a_ plots are down sampled). **B)** Frequency dependence of the average cumulative LFP response, N_2a_ and N_2b_ measured at 35s during stimulation at different frequencies in the vermis (n=10) and hemisphere (n=10) (same slices as in panel A). Statistical significance is indicated by asterisks (* p<0.05).

The LFP amplitude was considered as a proxy of NA. However, N_2a_ and N_2b_ showed an almost exponential decrease during the stimulus trains, mostly reflecting adaptation in the neurotransmission process ^53^, with time-constants that became smaller at higher frequencies. This trend markedly differed from that of NVC (cf. **Fig.1C**), so that neither N_2a_ nor N_2b_ nor their sum explained the time-course of dilation and its non-linearity in the frequency domain (**Fig. 4B**).

### Simulated NMDA currents correlate with NVC time course

The biochemical pathway leading to capillary dilation in the granular layer relies on activation of postsynaptic NMDARs in granule cells causing neuronal nitric oxide synthase (nNOS) activation and NO release ^21^. Since the NMDA LFP component could not be quantitatively extracted from N_2b_ (not being the only contributor to this peak), we used a realistic computational model of the cerebellar network ^46^ to simulate the NMDAR component of granular layer responses to mossy fiber inputs. The model of the vermis was the same used previously ^46, 47^, while the model of the hemisphere was adjusted to tune cell densities according to the Allen Brain Atlas (see Methods). Accordingly, granule cell density was increased resulting in a higher granule cell / Golgi cell ratio (**Fig.5A**). Model validation was performed against the experimental data at different time points and frequencies, showing a high level of convergence (**Fig.5B**; see also Methods for details). Eventually, the model yielded the NMDA current build-up in granule cells during the different stimulation patterns in the vermis and hemisphere.

**Fig. 5.**
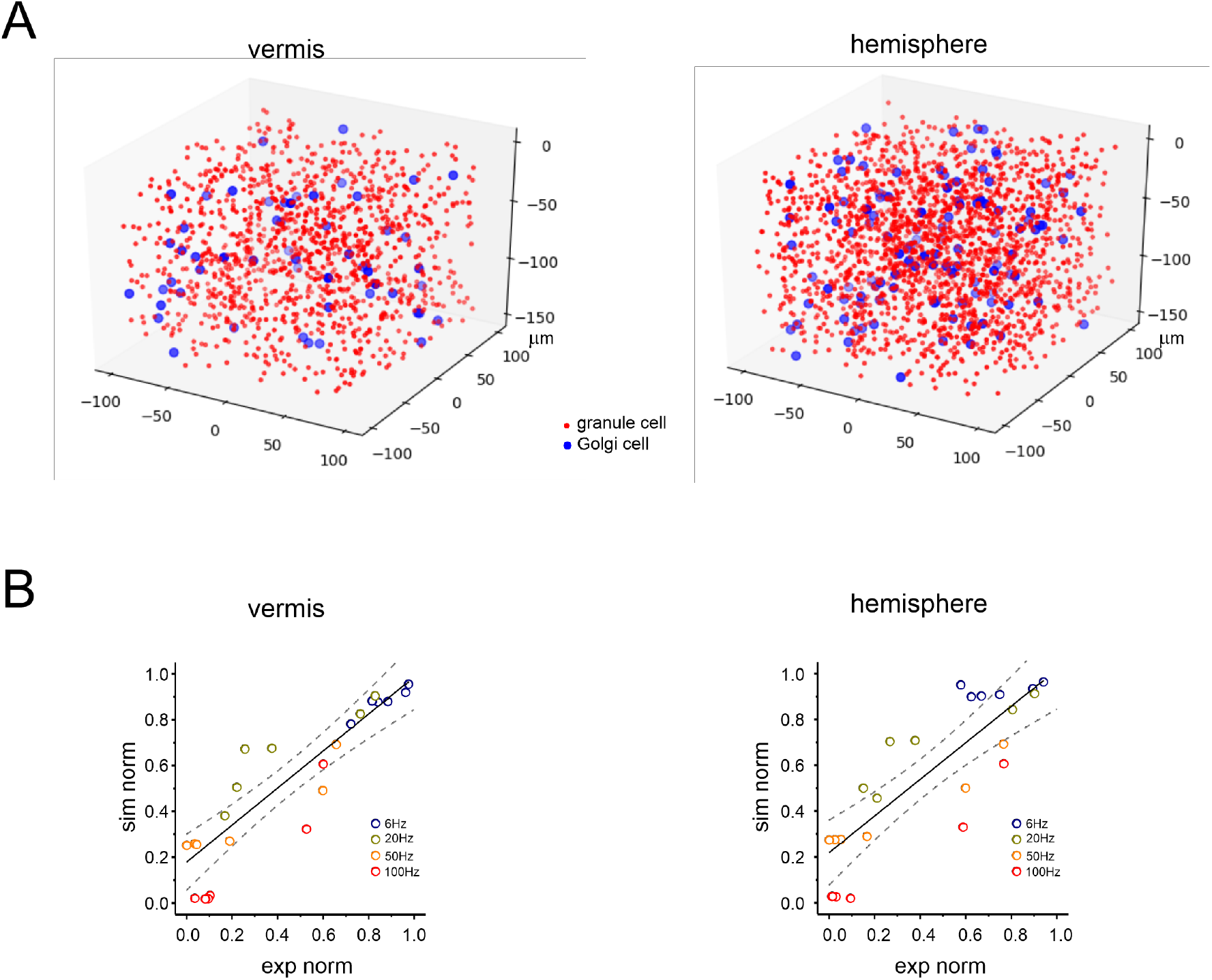
Realistic computational modeling of granular layer response in the vermis and hemisphere. **A)** Schematic illustration of the spatial distribution of granule cells (*red dots*) and Golgi cells (*blue dots*) in a volume chunk of the granular layer (100µm^3^) of the cerebellar vermis and hemisphere in a realistic computational model derived from ^46^. For better rendering, granule cells are down sampled by a factor of 20. There are 1203 granule cells and 55 Golgi cells for the vermis, 1980 granule cells and 113 Golgi cells for the hemisphere. **B)** Computational model validation against the experimental data for the vermis and hemisphere. The plot shows the distribution of normalized response amplitude experiments (N_2a_ peak amplitude) and simulations using the same stimulation frequency. The time points considered for model validation are 1s, 2s, 10s, 20s, 25s, 35s. The simulated frequencies are the same used in the experiments. The black lines are linear fittings (*vermis*: slope 0.81, R^2^ 0.73, p<0.001; *hemisphere*: slope 0.80; R^2^ 0.70, p<0.001), the dashed lines are confidence intervals.

We then considered that vasodilation in the granular layer is mediated by NMDAR-dependent production of NO ^21^ acting, through volume diffusion, on guanylyl cyclase (GC) in pericytes. The consequent cyclic guanosine monophosphate (cGMP) levels are the ultimate cause of vasodilation and are regulated by the enzyme phosphodiesterase (PDE) ^54^. We then simulated the cumulative NMDA current build up, corrected by PDE (−26.4% and −59.9% after 20s and 35s, respectively ^54^), and used it as a proxy of the signal controlling NVC (**Fig.6A**). Indeed, the corrected NMDA current build-up approached NVC, at all frequencies tested (**Fig.6B and Supplementary Fig.S1**).

**Fig. 6.**
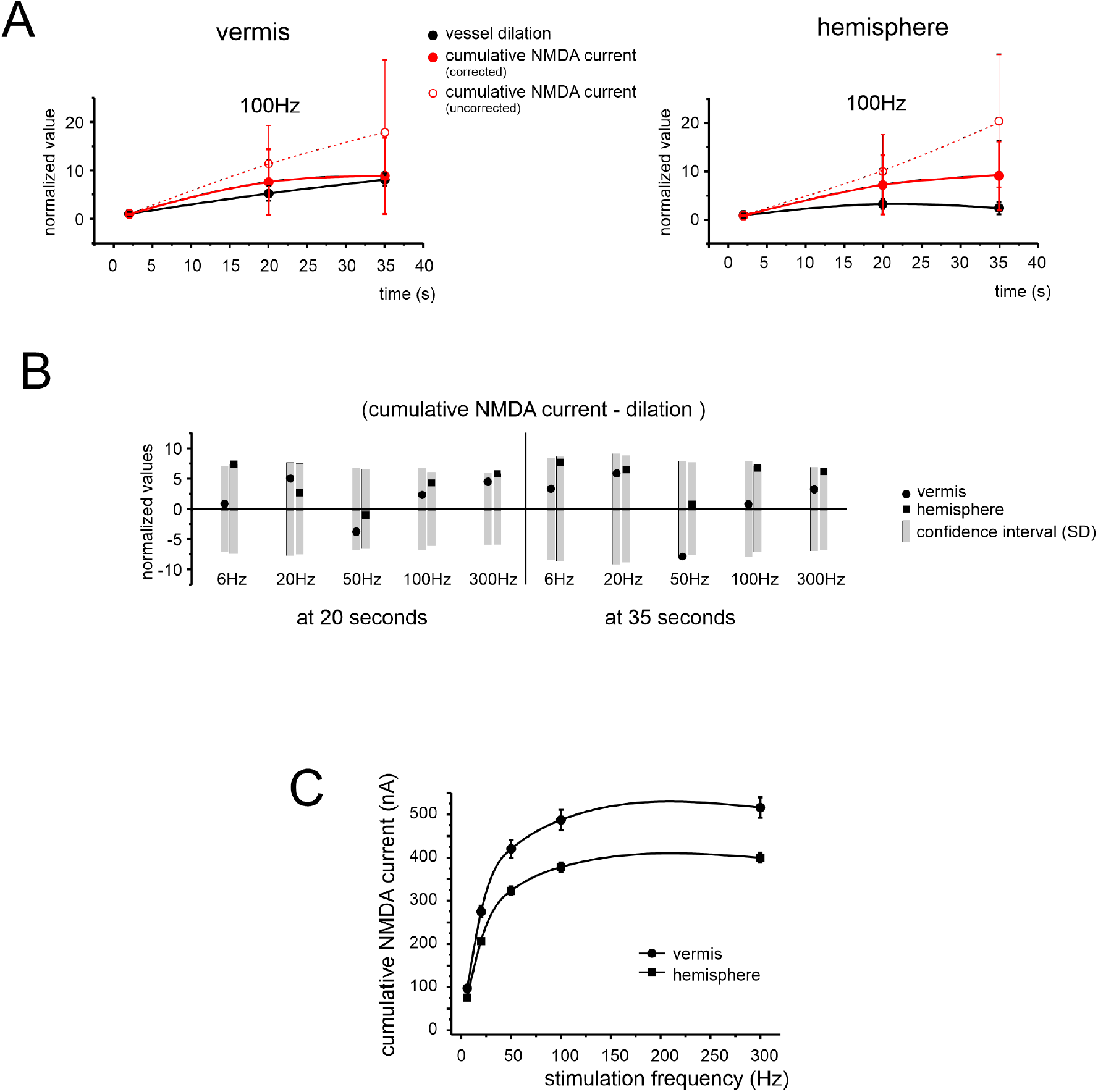
Comparison of simulated NMDA currents in granule cells and blood vessel changes. **A)** Time-course of the granule cells cumulative NMDA current during 100Hz stimulation and of the corresponding vessel response, for the vermis and hemisphere. The plot shows both the corrected and uncorrected NMDA component (see text for details). All data are normalized to amplitude at 2s. **B)** The bar graph shows the difference between the normalized values of simulated NMDA component and vessel dilation at 20s and 35s at all tested frequencies, for the vermis and hemisphere. The grey rectangles represent the normalized confidence interval (±1SD) yielded by model simulations of the NMDA current. Notice that all data points fall within the confidence interval, both for the vermis and hemisphere. **C)** Frequency-dependence of the cumulative NMDA currents measured at 35s, for the vermis and hemisphere. This panel should be compared to the corresponding one for vasodilation (cf. **Fig.2B)**.

In aggregate, these simulations suggest that the NMDAR-NO-GC pathway, incorporating the effect of PDE on cGMP levels, is sufficient to explain the time-course of vasodilation, at each frequency tested, both for vermis and hemisphere. Conversely, NMDAR activity alone was not sufficient to explain either the frequency-dependence or region-dependence of NVC (**Fig. 6C** to **Fig. 2B**).

## Discussion

The central finding in this study is that NVC in the cerebellum shows non-linear frequency-dependence. The capillary diameter increased in response to mossy fiber activation in the 6-300 Hz range, with a marked inflection around 50 Hz (vermis) and 100 Hz (hemisphere). The NMDA receptor-mediated component of granular layer responses triggering nNOS activation and NO production, once corrected for PDE activation (**Fig.7A**), could effectively explain the time-course of vessel dilation at each stimulus frequency but was insufficient to explain the non-linearity in vessel response with respect to input frequency. These results reveal complexity and diversity of NVC at the microscale suggesting a partial uncoupling from NA and the intervention of other mechanisms in addition to the NMDA-nNOS-NO pathway.

**Fig. 7.**
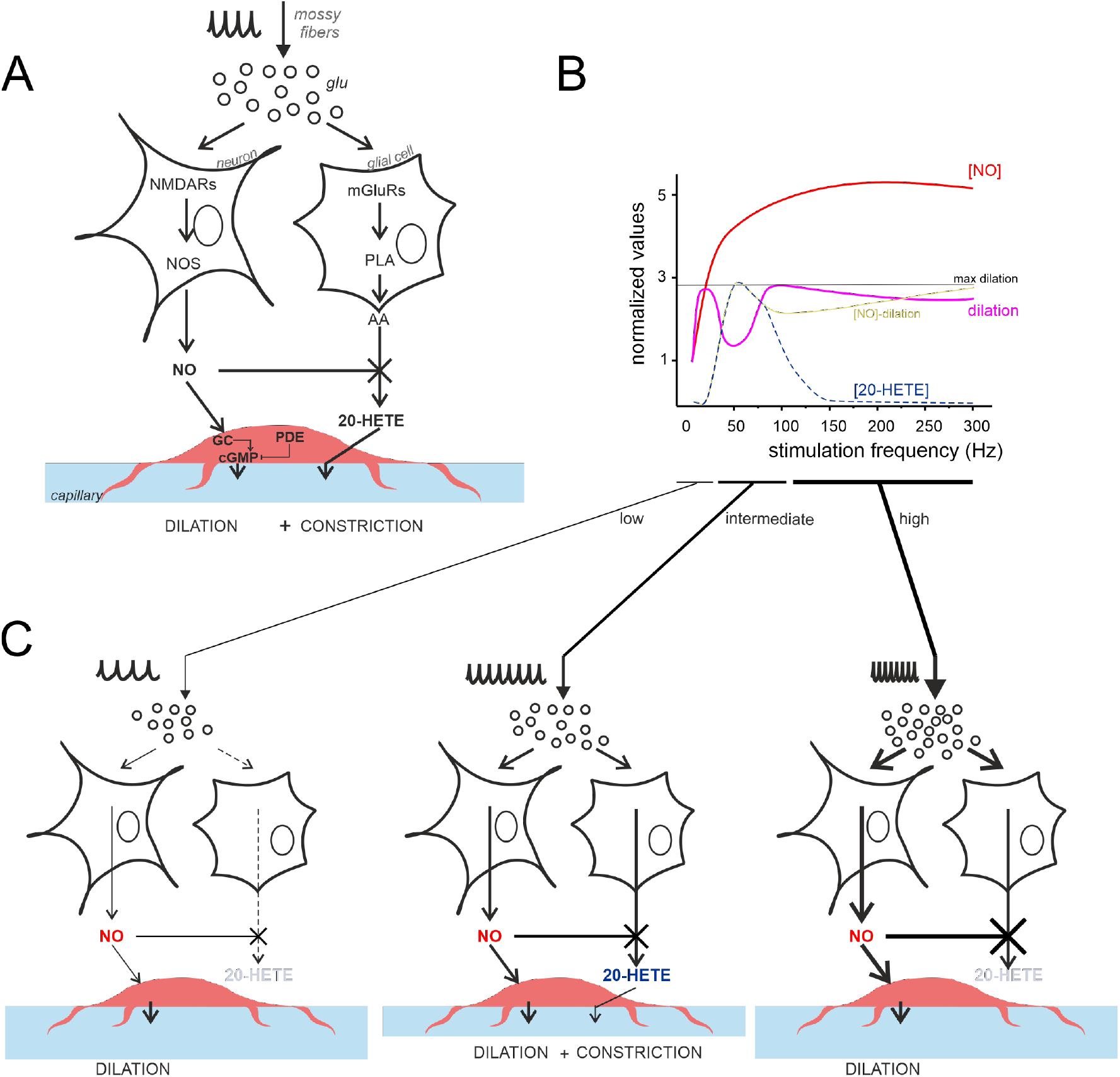
The *vasodilation-vasoconstriction competition* hypothesis. **A)** Schematic illustration of the main players involved in NO and 20-HETE release, according to ^21^. Vasodilation is mediated by the NMDAR-NOS-NO pathway while vasoconstriction is mediated by the mGluR-PLA-20-HETE pathway. Notice that NO inhibits 20-HETE synthesis ^8^. PLA, phospholipase A; GC, guanylyl cyclase; PDE, phosphodiesterase. **B)** Schematic illustration of the competition between NO and 20-HETE in determining vessel diameter changes. [NO] (*red*) is directly proportional to the simulated NMDA current (not shown), while capillary dilation (*violet*) is provided by experimental measurements (the difference between the two curves is in *yellow*). 20-HETE is specular to the NVC inflection at intermediate frequencies (more details are given in text). **C)** The differential engagement on the NO and 20-HETE pathways are shown for low, intermediate, and high stimulus frequencies (note the different thickness of the arrows).

### Non-linearity and region specificity of cerebellar NVC

Mossy fiber stimulations induced capillary vasodilation in the mouse cerebellar granular layer, attaining a maximum around 35 seconds. This effect, which was observed both in the vermis and hemisphere, resembles that previously reported in rat cerebellar slices ^21^. Interestingly, vasodilation reached the maximum already at 20Hz but then decreased to raise back to maximum at 300 Hz. These data demonstrate that cerebellar NVC is non-linear with respect to circuit stimulation frequency. The NVC curve, after bending beyond 20 Hz, recovered more rapidly in vermis than hemisphere. It is worth noting that capillary dilation ranging from ∼2.5% to ∼7.5% is sufficient to impact the total blood flow, which is directly proportional to the fourth power of the vessel radius according to Poiseuille’s law ^21, 55^.

The reasons of the region specificity of cerebellar granular layer NVC remain unclear. Both in the granular layer of cerebellar vermis and hemisphere, the pericytes were detected on capillary walls and in close contact with granule cells, i.e. in the ideal location to detect the chemical mediators released by neurons and glial cells in response to local NA changes ^22, 50^. Thus, the fundamental anatomical organization of the main cellular components was similar. Some differences in NA (relative size of the N_2a_ and N_2b_ LFP components) were observed between the two regions but did not explain those in NVC. Thus, neither anatomical nor electrophysiological data could explain local NVC differences. Therefore, there is a partial uncoupling of NA from NVC and metabolic effects may play a role along with the neurogenic ones ^7^. This case resembles that reported in the cerebral cortex, where differences in CBF occurred between the somatosensory and frontal areas of awake mice during locomotor activity on a treadmill ^56^ in the absence of differences in NA.

### NA and NVC in the cerebellar vermis and hemisphere

Reports in various brain areas have identified the ∑LFP as the measure of NA that best matches NVC ^28-36^. Here, we used a high-density multi-electrodes array (HD-MEA) to record LFPs from the granular layer of cerebellar cortical slices in the vermis and hemisphere during mossy fibers stimulation. The high spatial resolution of this technique allowed us to record from up to nine electrodes within 200µm from the stimulation site, bringing the precision of our measurement on scale with the inter-capillary distance. Both in the vermis and hemisphere, the LFP was composed by the typical N_2a_-N_2b_ wave sequence ^45, 51, 52^, which is informative on the AMPAR- and NMDAR-mediated components of granule cells responses to mossy fibers stimulation, respectively. Interestingly, N_2a_ was larger in the vermis while N_2b_ was larger in the hemisphere. These differences were well matched by updating the granule cell-Golgi cell ratio and neuronal density in the hemisphere and the vermis according to the Allen Brain Atlas in a realistic microcircuit model (see Methods for details) and could therefore reflect differences in local network organization. We expected that the differences in NA could explain the NVC non-linearity. However, the LFP parameters did not account for NVC dynamics neither in the time nor frequency domains.

### NMDAR currents drive the time-course of dilation but do not determine its frequency-dependence

Unlike other brain regions, capillary vasodilation in the cerebellar granular layer is known to entirely rely on the NMDAR-nNOS-NO pathway, in line with the fact that this layer has the highest level of nNOS expression in the brain ^14, 21^. Moreover, a previous report in the rat somatosensory cortex showed the failure of ∑LFP in accounting for CBF variations at specific frequencies when the NMDA currents did not contribute to the signal ^29^. It should not be surprising then that the parameter describing the relationship between NA and the time-course of dilation in our case is the NMDAR-mediated component of granule cells response. In this study, a realistic theoretical model of the granular layer, tuned against subtle anatomical differences between vermis and hemisphere, was crucial to reconstruct the NMDAR-mediated component of the response. Following validation against experimental data, the simulated NMDA current build-up proved suitable to determine the time-course of vasodilation. Nevertheless, the NMDA current could neither explain the frequency-dependence of vasodilation (see **Fig. 2B**) nor its regional specificity.

### The vasodilation-vasoconstriction competition hypothesis

In a previous study ^21^, we demonstrated that NVC in the granular layer is mediated by NA-dependent release of the vasodilator agent NO, which occludes the effect of the vasoconstrictor agent 20-HETE, released following mGluRs activation presumably in glial cells. The effect of 20-HETE appeared only when NOS activity was blocked, in line with the notion that NO blocks this vasoconstrictor pathway ^8^ (**Fig.7A)**. The inflection in the NVC-frequency plot hints for a competition between vasodilator and vasoconstrictor pathways based on the frequency-dependent balance between NO and 20-HETE production (**Fig.7B**). Indeed, in rat cerebellar slices, pharmacological subtraction experiments revealed that 20-HETE reduced by 44% the vasodilation mediated by NO at 50Hz, in fair agreement with the data reported here ^21^.

The *vasodilation-vasoconstriction competition* hypothesis is illustrated in (**Fig.7C)**. The amount of NO produced at low frequency (<50 Hz) is sufficient to block 20-HETE synthesis. Indeed, 20-HETE depends on the mGluRs activation, which needs neurotransmitter accumulation during high-frequency input discharges ^57^. At increasing frequencies, glutamate build-up and spillover would intensively activate extra-synaptic mGluRs ^57, 58^ recruiting the 20-HETE pathway. At this point, 20-HETE synthesis would surge, NO would not be able to counteract it, and the pericytes would integrate the two opposite signals causing the inflection of NVC curves observed at intermediate frequencies (50-150 Hz). At high frequency (>50-100 Hz), NO would increase enough to block the 20-HETE synthesis. This effect is probably helped by mGluRs desensitization and saturation of intracellular calcium concentration in glial cells at increasing glutamate concentrations ^59, 60^. Although not exhaustive, this set of mechanisms accounts for the NVC/frequency non-linearity and provides the basis for a future quantitative mathematical model. Needless to say that such model would provide a mean to understanding neurological and neurodegenerative diseases, such as multiple sclerosis, where the non-linear BOLD response to a variable grip force task was altered in the primary motor cortex, with a change in the non-linear behavior especially in patients with greater disability (Alahmadi 2021). Future work should aim not only to improve the mathematical model explaining the NVC complexity from cellular physiology experiments, but to scale such model to interpret *in vivo* data from human studies of central nervous system pathologies ^61^.

### Comparing different NVC hypotheses and the case for cerebellar region specificity

The *vasodilation-vasoconstriction balance*, different from an NA *ceiling* effect, could prevent the system from saturating through a dynamic regulation of the vessel caliber set-point when the input frequency increases. However, it does not account alone for the shift from 50Hz to 100Hz in curve inflexion between vermis and hemisphere and requires further comments. We cannot rule out that subtle anatomical differences occur beyond the resolution of this study, addressing the “neurogenic hypothesis” of NVC. Other factors that are likely to come into play have been reported in different brain regions. Activity-dependent metabolites and lowering of glucose and oxygen concentrations can link neuronal activity and vessel tone (e.g. see discussion in ^7^) addressing the “metabolic hypothesis”. It is also interesting to consider the “set-point hypothesis” for determining dilation or constriction depending on the initial vascular tone ^11^. Although we did not find any difference in pre-constricted capillary diameter between vermis and hemisphere, we cannot rule out the possibility that different set-points characterize these two regions, so that similar initial diameter would not imply the same dilation/constriction balance. In addition, signal integrating cells could include not just the pericytes but also astrocytes ^11^ and endothelium ^12^, although blood vessel intrinsic properties should have been marginally involved due to the lack of intravascular pressure in acute brain slices. In summary, it is highly probable that the NVC does not simply rely on the metabolic or neurogenic hypotheses but is defined by a complex interaction of multiple factors that vary locally from one brain region to another.

### Conclusions

The mechanisms of NVC and its impact on local brain functioning are raising increasing attention in the scientific community, both for understanding BOLD signals used in fMRI and for the potential implications in neurovascular pathology. Our data show that NVC is probably more complex than previously thought, adjusting to the input frequency in a region-specific manner. To the best of our knowledge, this is the first report of this effect in the cerebellar cortex. Interestingly, our results suggest that the force/BOLD non-linearity recorded from the cerebellum during motor task execution ^24, 25^ could, at least in part, be due to local non-linear NVC properties. Although the understanding of NVC finetuning is still incomplete, our results integrate the neurogenic and metabolic hypothesis opening the perspective of a dynamic microvessel diameter regulation, which should be considered to interpret BOLD signals in fMRI recordings.

## Acknowledgements

This research has received funding from the European Union’s Horizon 2020 Framework Programme for Research and Innovation under the Specific Grant Agreement No. 945539 (Human Brain Project SGA3) and Specific Grant Agreement No. 785907 (Human Brain Project SGA2) and from Centro Fermi project “Local Neuronal Microcircuits” (LNM) to ED. CGWK receives funding for research activities from the MS Society (#77), Wings for Life (#169111), BRC (#BRC704/CAP/CGW), UCL Global Challenges Research Fund (GCRF), MRC (#MR/S026088/1), Ataxia UK.

## Author contributions

GG performed and analyzed microvessel recordings and confocal images; GG and AM performed and analyzed electrophysiological recordings; ED designed the model and SC performed the simulations; UL performed double immunofluorescence; CGWK provided the initial question and critical revision of the intellectual content; LM analyzed the data, wrote the manuscript, and prepared the figures; LM and ED designed the project and coordinated the work. ED supported it financially and finalized the manuscript.

## Conflict of interest

The authors declare that the research was conducted in the absence of any commercial or financial relationships that could be construed as a potential conflict of interest.

## Data availability

All the materials related to the paper are available from the corresponding authors upon reasonable request.

## Notes

### Competing Interest Statement

The authors have declared no competing interest.

