## supplemental materials for "Non-linear frequency-dependence of neurovascular coupling in the cerebellar cortex implies vasodilation-vasoconstriction competition"

### Supplementary figure legend

**Fig. S1 – Model-derived NMDA component of granule cell responses: comparison to vessel response at different stimulation frequencies.** Time-course of the granule cells cumulative NMDA current (*red*) building-up during the simulation at different frequencies and of the vessel responses (*black*) in the corresponding conditions. Both parameters are normalized for their amplitude at 2s, to compare the trends over the stimulation time. Notice that the trends of the two parameters do not significantly differ at any frequency tested, both for the vermis (*at left*) and hemisphere (*at right*).

**Table 1 – Maximal capillary dilations at different stimulation frequencies.**

The table reports the average maximal dilations (percent changes compared to the baseline) observed at different stimulation frequencies in the vermis and hemisphere (paired Student's t test: *p<0.05; **p<0.01; ***p<0.001).

| | 6Hz | 20Hz | 50Hz | 100Hz | 300Hz |
|---|---|---|---|---|---|
| **vermis** | 2.99±0.61(***) | 7.19±1.85(***) | 3.66±0.82(***) | 7.24±1.26(***) | 6.42±1.10(***) |
| **hemisphere** | 4.93±1.06(**) | 6.96±1.45(***) | 5.79±0.91(***) | 4.28±1.00(***) | 7.04±1.28(***) |

**Table 2 – N2a and N2b peak amplitude changes at the end of the stimulations.**

The table reports the average percent changes of N2a and N2b peak amplitudes comparing the last response to the first one in the stimulation train, at the different frequencies used (paired Student's t test: *p<0.05; **p<0.01; ***p<0.001).

| | 6Hz | 20Hz | 50Hz | 100Hz |
|---|---|---|---|---|
| **vermis N2a** | -20.3 ± 4.3 (*) | -41.7 ± 4.4 (***) | -52.4 ± 3.9 (***) | -52.2 ± 3.4 (***) |
| **vermis N2b** | -3.0 ± 4.3 (**) | -12.3 ± 4.5 (***) | -7.2 ±5.6 (***) | -2.8 ± 6.3 (***) |
| **hemisphere N2a** | -22.1 ± 4.4 (***) | -47.9 ± 3.9 (***) | -54.9 ± 2.5 (***) | -54.7 ± 2.8 (***) |
| **hemisphere N2b** | -23.1 ± 6.4 (***) | -22.8 ± 6.2 (***) | -29.1 ± 5.5 (***) | -23.0 ± 5.5 (***) |
