## Supplementary figures and images for "Non-linear frequency-dependence of neurovascular coupling in the cerebellar cortex implies vasodilation-vasoconstriction competition"

### Figure S1

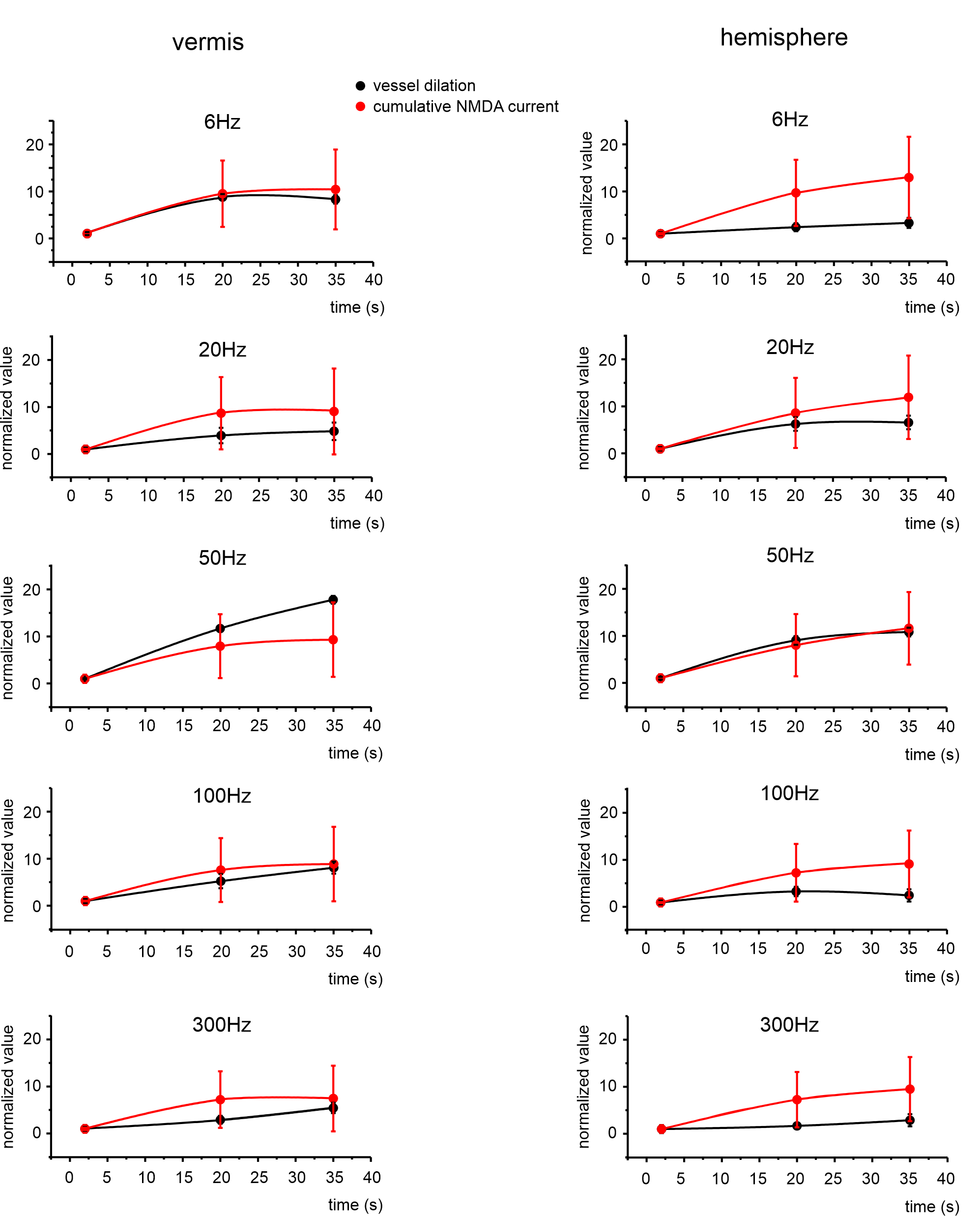
